# Microscopic Dissipative Structuring at the Origin of Life

**DOI:** 10.1101/179382

**Authors:** Karo Michaelian

## Abstract

Fundamental molecules of life are suggested to be formed, proliferated, and evolved through microscopic dissipative structuring and autocatalytic replication under the UV-C solar spectrum prevalent at Earth’s surface throughout the Archean. Evidence is given in the numerous salient characteristics of these, including their strong absorption in this spectral region, their rapid non-radiative decay through an inherent conical intersection, UV-C activation (phos-phorylation) of nucleotides, and UV-C induced denaturing of double helix RNA and DNA. The examples of the dissipative structuring and dissipative proliferation of the purines and of single strand DNA are given. This provides a physical-chemical foundation for understanding the origin and evolution of life.

## 1. Introduction

It is well known empirically, and described theoretically within the framework of Non-linear Classical Irreversible Thermodynamics (Prigogine, 1967), that “spontaneous” macroscopic organization of material in space and time can occur within a system subjected to an externally imposed generalized chemical potential. This happens as the result of a non-equilibrium thermodynamic imperative to increase the dissipation of the external potential, i.e. to produce entropy, or, in still more fundamental terms, to spread the conserved quantities of Nature (energy, momentum, angular momentum, charge, etc.) over ever more microscopic degrees of freedom. This phenomenon of the “spontaneous” structuring of material under a generalized chemical potential is known as *dissipative structuring*. Prigogine was awarded the Nobel Prize in chemistry in 1977 *"for his contributions to non-equilibrium thermodynamics, particularly the theory of dissipative structures"* (Nobel committee, 1977). Earlier, however, Boltzmann (1886) had realized that material was organizing to dissipate the impressed solar photon potential and Schrödinger (1944) emphasized this fact and brought Boltzmann’s deep insight into the forefront of scientific investigation on the physics and chemistry of life.

For any system maintained out of equilibrium through the imposition of an external generalized chemical potential, the existence of non-linearity between the induced generalized flows and forces derived from the potential will endow the system with numerous possible solutions for the time-relaxed non-equilibrium stationary state. These solutions often correspond to stable dynamic material organizations or processes of low entropy, involving symmetry breaking in both space and time. As will be illustrated below, which of the possible stable solutions the system evolves into will depend on the initial conditions and on a particular random microscopic fluctuation at a bifurcation point which gets amplified into the solution. This endows the evolutive process with a particular history.

An instructive example of dissipative structuring occurring in a system held out of equilibrium by an external generalized chemical potential is that of the formation of Bénard cells (see figure 1). These convection cells arise in a liquid system held under a temperature gradient maintained by a lower reservoir held at high temperature and an upper reservoir held at a lower temperature, the whole system being subjected to a gravitational field. At a sufficiently large temperature gradient, at which the system is forced into the non-linear regime between heat flow and the force corresponding to the temperature gradient, a particular microscopic fluctuation that produces a small movement of a particular heated volume element of the fluid towards the upper plate will be reinforced because of its lower density with respect to the water above it. If the fluctuation is large enough, then this buoyant force for the upward movement of the volume element of the fluid will overcome the viscous frictional forces acting against the movement and over the conduction of heat which tends to dampen macroscopic dynamics within the system, and the volume element will continue on its upward trajectory, pulling in warmer liquid behind it. In this manner a convection cell, known as a Bénard cell, is established. This cell, whose dimension depends on the physical characteristics of the solvent material, the confining volume, and the imposed potential, will now provide a strong local perturbation, stimulating neighboring volumes of liquid to also form cells, and very quickly the whole system fills with Bénard convection cells. The initial cell, or, in fact, a small macroscopic portion thereof, acts as a catalyst for a second similar cell, and so on in an exponential manner, all started by a single random fluctuation of a microscopic volume element at the bifurcation point.

**Fig. 1:**
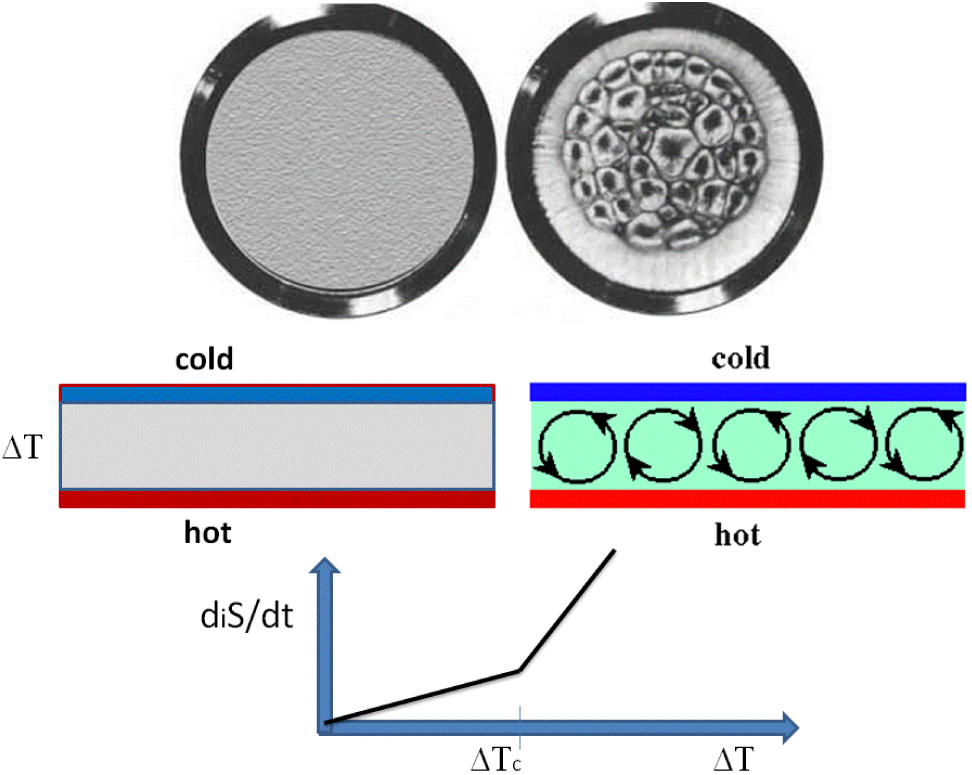
Bénard Cells: The onset of structuring of material in space and time as a result of an external potential imposed over the system (in this case a temperature difference) and a non-linear relation between flows (heat flow) and forces (temperature gradient). The left part of the top panel describes the linear situation for the relation between the flow and the force when the temperature difference over the system is below some critical value ΔT_c_ and the system remains homogeneous (no spatial symmetry breaking) and there exists only conduction of heat (middle panel left). The right part of the top panel describes the non-linear situation when the temperature difference between the two plates is greater than some critical value ΔT_c_ for which convection cells spontaneously arise (Bénard cells) and spatial symmetry breaking occurs (middle panel right). The direction of liquid circulation in the convection cell is determined by an initial microscopic fluctuation at the bifurcation point corresponding to ΔT_c_ which grew into the cell. Two solutions are allowed for cells at the bifurcation point, one with clockwise rotation and the other anti-clockwise. In the lower panel the internal entropy production is plotted as a function of the temperature difference over the system. A further bifurcation into two stationary state solutions with distinct entropy production is also possible; that of the hot solvent rising in the center of the cell (the greater entropy producing and more stable solution) or hot solvent rising at the edges of the cell (the lower entropy producing, less stable solution). There is an increase in slope of the entropy production as a function of temperature gradient at the critical value of the temperature difference, ΔT_c_, required for the onset of convection. Image credit: Adapted, Public Domain.

Is there a lower limit to the size to these Bénard structures? In a Bénard convection system, there are two irreversible processes occurring; heat conduction and convection. Convection spontaneously arises when the physical conditions allow it, when the upward buoyant force due to the lower density of a heated liquid volume element becomes greater than the viscous forces of friction that oppose its movement. This requires a sufficient temperature gradient to be maintained over the system, and the conduction of heat to be sufficiently slow from the hot plate to the cold plate so that convection remains competitive with conduction. As the distance between the two plates is decreased, heat diffusion (conduction) becomes ever more "instantaneous" and even though the temperature gradient becomes greater, Bénard convection cells eventually cannot compete with conduction at heat dissipation. At these small dimensions, approximately ten’s of microns, only conduction remains and structuring disappears.

As another example of a lower limit to the size of a dissipative structure, consider a macroscopic reaction-diffusion system, such as that giving rise to the Belousov-Zhabotinsky chemical oscillator (see figure 2). In this system there are also two irreversible processes occurring; diffusion and chemical reactions. The characteristic diffusion times (the average time between scattering events) are very large compared with typical chemical reaction times and so reactions can be considered to occur instantaneously. Alan Turing (1952) predicted that in such a system of reacting and diffusing chemicals, macroscopic patterns of products and reactants would spontaneously emerge, breaking homogeneity in space and time, if there existed an autocatalytic agent (an activator) who's efficacy was limited by an inhibitor, thereby introducing non-linearity into the system through feedback. In a reaction-diffusion system, the characteristic size of the macroscopic pattern is, by necessity, greater than the average diffusion length. For biological systems in a water solvent environment, the minimum size for dissipative structuring of this reaction-diffusion type is also of the order of microns (Hess and Mikhailov, 1994).

**Fig. 2:**
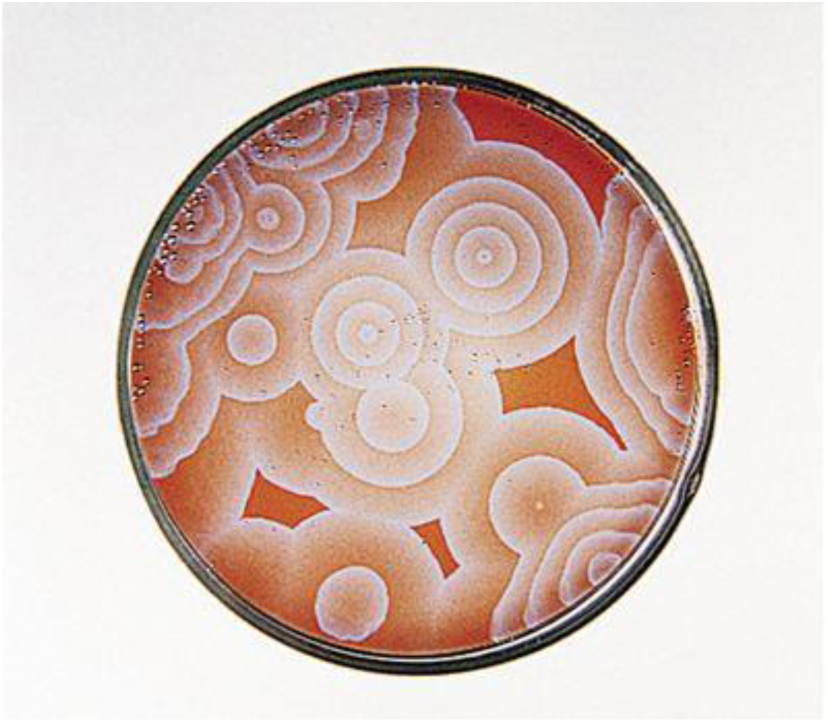
Turing patterns of concentration gradients of reactants and products formed in a Beluosov-Zhabotinsky system of chemical reactions with diffusion. With a constant source and sink for the reactants and products (i.e. with an imposed constant chemical potential defined by the environment) the patterns oscillate in space and time giving the appearance of a living system. This breaking of space and time symmetry is a non-equilibrium thermodynamic result and it is driven by the thermodynamic principle of increasing the rate of global entropy production. Turing patterns have a lower limit in size of about a few microns. Image credit: Public Domain.

Therefore, a Bénard dissipative conduction-convection, or a Belousov-Zhabotinsky reaction-diffusion, process could not give rise to structuring at the sub-cellular scale. However, dissipative structuring at the sub-micron, and even at the nanometer, scale occurs as witnessed by the many processes occurring inside a living cell. An example being molecular motors, such as those promoting the advance of polymerase during the extension of RNA or DNA (Bustamante et al., 2005), or the process of the formation of peptide bonds between selected amino acids, known as translation, which occurs in organelles of the cell known as the ribosomes. There must, therefore, be other dissipative mechanisms besides convection-conduction and reaction-diffusion operating at the nanoscale level which drive these microscopic irreversible processes and their associated microscopic dissipative structuring.

At the nanoscale dimension, particularly for soft (biological) material there is cohesion between atoms due to strong and directional covalent bonds and this allows for molecular structural transitions; for example between different isomeric, tautomeric, or charged states of the molecule. There is also the possibility of transitions between the different electronic, vibrational, or rotational states, between different orientations of electric or magnetic dipole moments under an externally imposed electric or magnetic field, or between molecular complexes or excimer states, or microscopic phase separation (gas-liquid-solid). These transformations are related to the *internal* degrees of freedom of the molecule, or the molecular complex, which are also available for the distribution of the conserved thermodynamic quantities such as energy, angular momentum, charge, etc.

Photon exchange and absorption processes in material can lead to quantum resonances which, through their probabilistic nature of decay, can lead to the breaking of classical reversibility (Prigogine and Stengers, 1997). The “footprint” of this breaking of reversibility depends on the amount of energy in the photon and the absorbing atoms involved (Lucia, 2017). Photon induced processes are also often associated with microscopic dissipative structuring since photons can deliver an intense amount of energy of low entropy locally on a very small amount of material. This leads to very large generalized forces at microscopic dimensions which can lead to microscopic self-organization as a response to dissipate the applied photon potential. A typical example of this is a lasing system. In a system operating in the lasing mode, photons of a particular chamber resonant wavelength stimulate atoms in the material to emit photons of the same wavelength in a coherent manner, forming a “self-organized” system of atoms emitting coherently from a particular excited state, and giving rise to a narrow beam of coherent mono-chromatic photons. Even though the organized state of the coherent output photon beam is of low entropy (as compared to an equilibrium black-body spectrum emitting the same amount of energy) global entropy production is positive since by vacating the lower lying vibrational states superimposed on the electronic excited state of the material, internal structuring allows the pumping field to be dissipated more efficiently than would happen without stimulated emission. A laser heats up under operation and it is this heat emitted into the environment that is a measure of the inherent net positive entropy production of the system.

Chemical or photochemical reactions or diffusion, together with the possible internal molecular structural transitions mentioned above, can lead to microscopic structuring through the dissipation of a generalized chemical potential. This new field of research, referred to as *microscopic dissipative structuring*, although already treated formally by Prigogine (1967) through the analysis of internal degrees of freedom (see below), is only recently receiving a lot of attention due to its potential for application in nanotechnology since dissipation induced nanoscale structuring can persist even after removal of the external generalized chemical potential, due to atomic and molecular mobility issues resulting from strong inter-atomic interactions, and can thus be used in a large number of novel practical applications where controlled nanostructuring is required. For example, Hess and Mikhailov (1994) have studied particular mechanisms for soft matter nanostructuring based on equilibrium phase separation of two molecular species at a surface and a photon-induced de-absorption process. Another example is the dissipative structuring of gold nanowires under UV photoreduction of of linear self-assemblies formed from Au(OH)_4^-^_ and tetra-alkyl ammonium ions at a water–chloroform interface (Soejima et al., 2009; 2014).

## 2. Thermodynamic Formalism for Treating Microscopic Dissipative Structuring

The non-equilibrium thermodynamic formalism for treating dissipation over internal degrees of freedom with microscopic flows and forces giving rise to microscopic structuring is similar to that for treating macroscopic flows and forces giving rise to macroscopic structuring and was first studied by Prigogine (1967). The starting point is the Gibb’s equation,

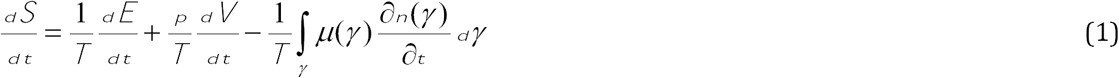

where n(*γ)* is the number of molecules in a state defined by some internal coordinate γ (for example the angle *ϑ* of the electrical dipole moment of the molecule with respect to an externally applied electric field), so *γ γ* is the number of molecules for which the internal coordinate lies between γ + *dγ* If we now first assume that the change of γ is discrete, i.e. γ can be changed by transformations from, or into, neighboring states γ -1 or γ +1, then the continuity equation is,

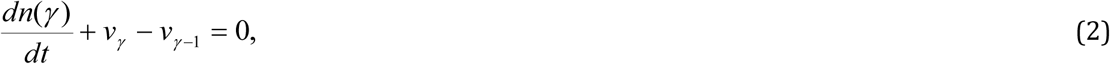

where *v*_*γ*_ is the rate for the transition *γ → γ* and *v*_*γ-1*_ is the rate for *γ → γ*. If γ isa continuous parameter, then the equivalent equation is,

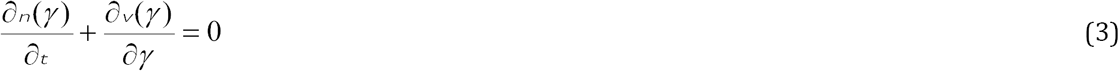

which is a continuity equation in the internal coordinate γ. *v*(*γ*) is the reaction rate (flow) giving the net flow of molecules into and out of the state defined by the coordinate γ. In vector notation, equation (3) becomes

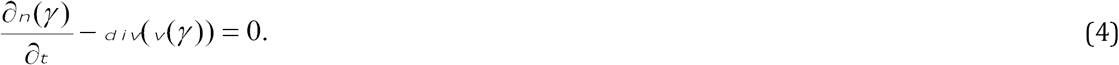

By partial integration, equation (1) can be transformed into

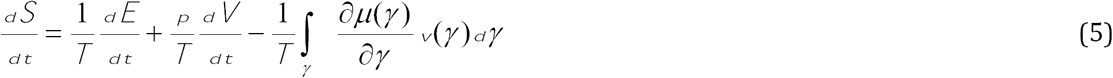

so the internal production of entropy due to the flow along the internal coordinate γ is

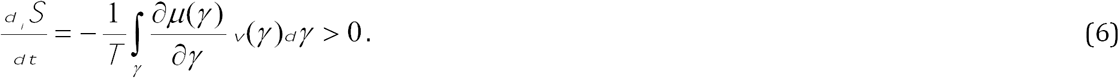

One can now postulate a further refinement of the second law of thermodynamics, valid within the microscopic internal coordinate space; *in each part of the internal coordinate space, the irreversible processes proceed in a direction such that a positive entropy production results*. This implies

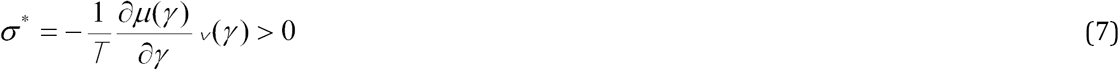

where *σ*_*_ is the entropy production per unit volume of the internal coordinate (or configuration) space. *σ*_*_ has the usual form of a product of an affinity or force, 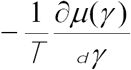, and a flow or rate, *v*(*γ*), of an irreversible process.

Just as in macroscopic non-equilibrium thermodynamics, linear relations between the forces and flows lead to a unique stationary state with a minimum of entropy production (with respect to the organization of the free forces *X*_*α*_ or flows *J*_*α*_; those not fixed by the environment). However, for non-linear relations between forces and flows, multiple stationary states exist, each with a possible different entropy production rate. Which solution is chosen by nature is determined by a microscopic fluctuation at a bifurcation, but it is more probable that fluctuations leading the system to greater entropy production will dominate the history of evolution towards the stationary state (Michaelian, 2016).

In the non-linear regime, the evolution of the system is no longer dictated by the principle of minimum entropy production, but rather by the *general evolutionary criterion* of Glansdorff and Prigogine (1964) which states that the variation of the entropy production, *P* = *d*_*i*_ *S/dt* = Σ_*α*_ *X*_*α*_ *J*_*α*_, due to variation of the free forces is negative semi-definite,

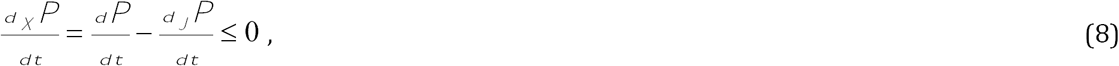

where *d*_*X*_*P/dt* is the variation of the entropy production with respect to the free forces and *d*_*J*_*P /dt* is the variation of the entropy production with respect to the free flows. In the stationary state *d*_*X*_*P/dt* = 0 and this criterion can be used to show (Prigogine, 1967; Michaleian, 2013) that, in the stationary state: 1) any product of a photochemical reaction (for example, a pigment), which acts as a catalyst for the dissipation of the same photon potential that produced it, will increase its concentration many-fold over what would be expected if the system were close to equilibrium or if the product did not act as a catalyst for dissipation, and 2) that the entropy production of the system increases. Given a constant external photon flux and a constant sink of the heat of dissipation, and the dispersal of the photochemical reaction products, this will lead to a continual proliferation of photochemical reaction products (pigments). Associating in this manner the fundamental molecules of life with autocatalytic microscopic dissipative structures (pigments), thereby explains the “vitality” of life at life’s beginnings and, indeed, throughout its evolutive history (Michaelian, 2016). Organisms are thus likely selected by nature not on the basis of some indefinable “fitness function”, but rather on the basis of their ability to catalyze dissipation and thereby secure their proliferation through this thermodynamic dissipation-replication relation.

An analysis of the particular photochemical reaction pathways involved in forming the most important fundamental molecules of life and the autocatalytic proliferation of these is given in the following sections after giving the evidence suggesting that they are microscopic dissipative structures.

## 3. The Fundamental Molecules of Life are Microscopic Dissipative Structures

Given the ubiquity of high energy photons arriving at the surface of planets, and the large number of photochemical reaction pathways available compared with thermal reactions starting from the electronic ground state, one would expect microscopic dissipative structuring to be common place. The fundamental molecules of life (those molecules found in all three domains of life) and their associations in polymers or other complexes are examples of microscopic dissipative structuring which occurred under the Archean UV-C photon potential (Michaelian, 2009; 2011; 2013; Michaelian and Simeonov, 2015; 2017).

Evidence for this is found in numerous characteristics of these, including the strong absorption cross sections, of almost all of the fundamental molecules, within the atmospheric UV-C window for light that was arriving at Earth’s surface during the Archean (Sagan, 1973, see figure 3). Many fundamental molecules also have conical intersections, or have chemical affinity to those fundamental molecules having a conical intersection, allowing donor-quencher energy transfer for rapid dissipation of the absorbed photon energy into heat.

**Fig. 3:**
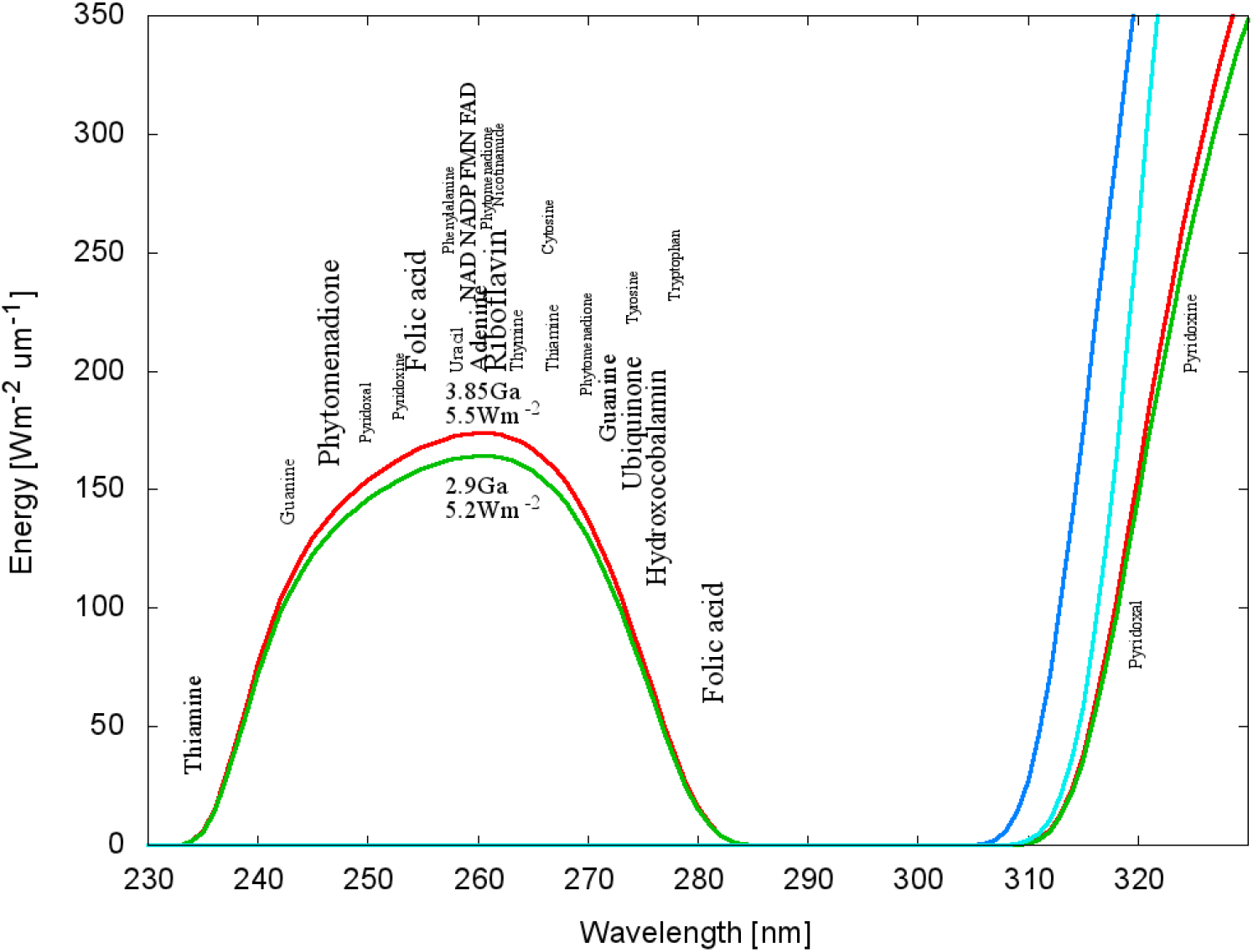
The wavelengths of maximum absorption of many of the fundamental molecules of life (common to all three domains) coincide with a predicted atmospheric window in the UV-C at the time of the origin of life at 3.85 Ga and until at least 2.9 Ga (red and green curves respectively). By 2.2 Ga (light blue curve) the UV-C light at Earth’s surface had been extinguished by the oxygen and ozone resulting from organisms performing oxygenic photosynthesis. The dark blue curve corresponds to the present day surface spectrum. The font size of the letter roughly indicates the relative size of the molar extinction coefficient of the indicated fundamental molecule (pigment). Image credit; adapted from (Michaelian and Simeonov, 2015).

The ubiquity of microscopic dissipative structuring of pigments in nature is also evidenced, for example, in the large quantity of organic pigment molecules found throughout the cosmos where UV light is available (Michaelian and Simeonov, 2017). Today, a similar situation exists on Earth with pigments dissipating in the near UV and visible regions which are self-organized through more complex biosynthetic pathways, but their production, nevertheless, remains as microscopic dissipative structuring employing the same light for their construction as they dissipate.

The nucleobases, as well their polymerization into single and double strand RNA and DNA, absorb and dissipate UV-C light. The nucleobases can be formed from simpler precursor molecules such as hydrogen cyanide HCN by the very same UV-C photons that they dissipate with such efficacy. The photochemical reaction pathways required for constructing many fundamental molecules still remain to be discovered, however, particular instructive examples of the dissipative structuring of the purines and of single strand RNA or DNA is given in the following sections.

## 4. Microscopic Dissipative Structuring of the Purines from HCN under UV-C Light

The most abundant carbon-containing 3-atom molecule in the observable universe is hydrogen cyanide HCN. Oró and Kimball were the first to discover a reaction pathway from HCN to the purine nucleobases (Oró, 1961; Oró and Kimballl, 1962). Later, Saladino et al. (2005; 2007) showed that all the nucleobases except guanine could be formed from formamide H_2_NCOH (a common product of HCN and H_2_O) in a water environment at high temperatures, of between 100 °C and 160 °C, with a number of distinct prebiotically available minerals acting as catalysts. However, more important to the thesis presented here, Ferris and Orgel (1966) discovered a low temperature generic photochemical pathway to the bases from HCN as given by the reaction scheme presented in figure 4. Finally, Barks et al. (2010) have shown that guanine could also be derived from formamide under UV irradiation at low temperatures.

**Fig. 4:**
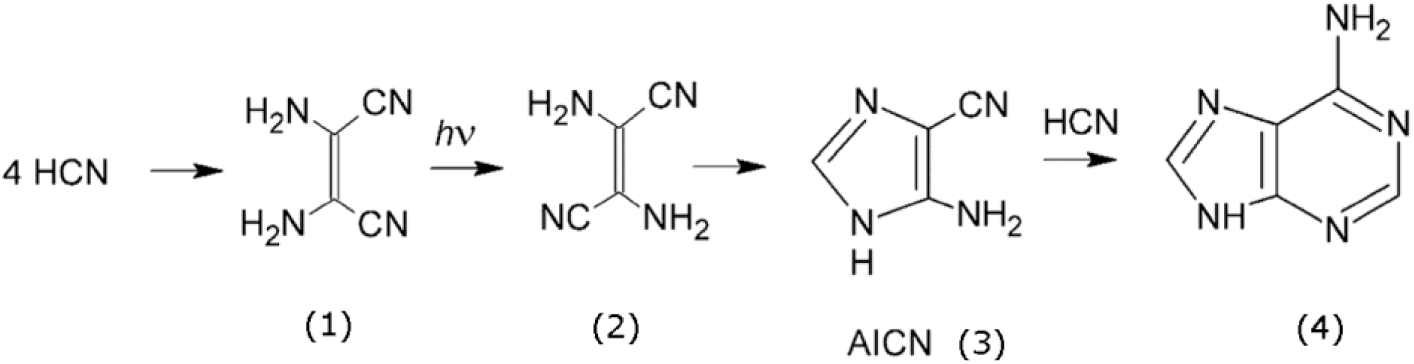
Generic photochemical pathway to the nucleobases first discovered by Ferris and Orgel (1966). Four molecules of HCN are transformed into the smallest stable oligomer (tetramer) of HCN, known as cis-2,3-diaminomaleonitrile (cis-DAMN), (1), which, under a constant UV-C photon flux isomerizes into the trans-DAMN (2) (also called diaminofumaronitrile DAFN) which may be further converted into an imidazole intermediate, 4-amino-1H-imidazole-5-carbonitrile, AICN, (3) and which together with another HCN molecule forms a purine (in this case, adenine), (4). Image credit: Boulanger et al., (2013), adapted from Ferris and Orgel (1966).

Photon disassociation of HCN is observed only for wavelengths < 200 nm (Cicerone and Zellner, 1983) so that once formed photo-chemically from N_2_ and CH_4_ (Trainer et al., 2012) in an oxygen-less upper atmosphere of early Earth and settling to the surface, it could accumulate without appreciable sinks (see figure 3).

The formation of the nucelobases at high temperature from HCN or formamide and inorganic catalysts as discovered by Oró (1961) and Oró and Kimball (1962) and further investigated by Saladino et al., (2005; 2007) may have been operative well before the origin of life at 3.85 Ga, perhaps even during the Hadean (4.6 - 4.0 Ga) when surface temperatures were well above 100 °C. However, by the origin of life at 3.85 Ga there is evidence that Earth’s surface temperature had cooled to ∼80 °C (Knauth, 1992; Knauth and Lowe, 2003), too cold for normal thermal chemistry with HCN and formamide, and the photochemical pathway to producing the nucleobases appears to be the most viable option so far discovered (Fig. 4). A more important reason for emphasizing this photochemical reaction pathway above other routes, however, is that this pathway exemplifies the characteristics of microscopic dissipative structuring and autocatalytic proliferation, which appear to be the relevant hallmarks of life even today. In the following, we therefore concentrate on the generic photochemical pathway to the purines as depicted in figure 4.

The actual processes leading from product (1) to (2) and (2) to (3) of figure 4 have recently been determined by Boulanger et al. (2013) and Szabla et al. (2014) by analyzing the possible mechanisms of the photochemical steps in detail using chemical kinetics and computational chemistry, employing density functional theory and time dependent density functional theory to determine the minima and transition states of the electronic ground state and excited state, respectively. The fast decay of the excess energy due to photon absorption and internal conversion into vibrational energy, with approximately 2/3 of the excess energy being dissipated within 0.2 ps to the surrounding water molecules, indicated that maximum free energy barrier heights for hot ground state thermal reactions are approximately 30 Kcal/mol (Boulanger et al., 2013). Any barriers higher than this would have to be overcome by involving a photon. Using such kinetic constraints and transition state formalism, they find that there is only one sequence of steps that is thermodynamically and kinetically compatible with experiment. This sequence is represented schematically in figure 5.

**Fig. 5:**
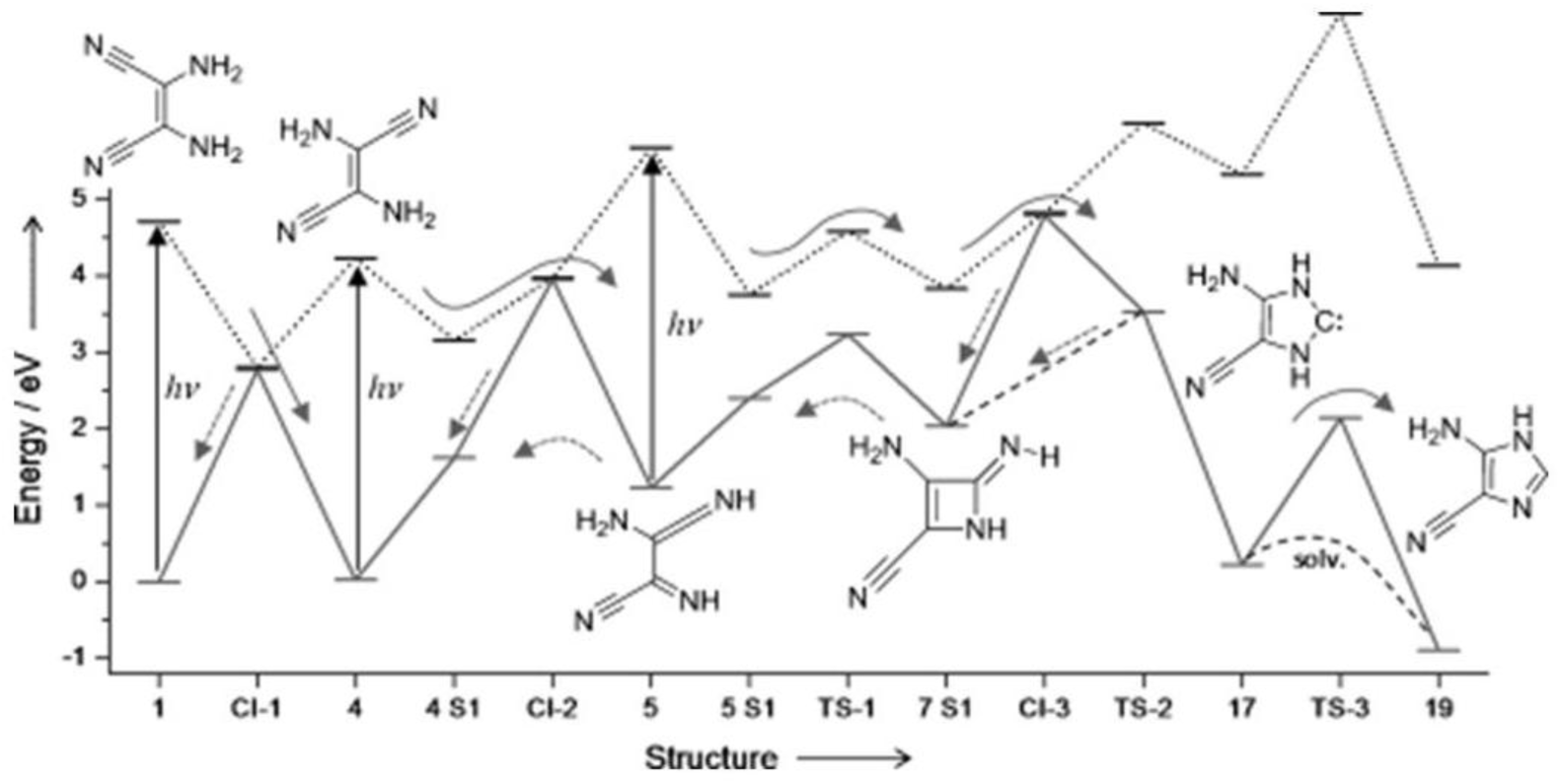
The sequence of photochemical and chemical reactions that lead from cis-DAMN at (1) to AICN at (19). The solid arrows correspond to the forward reactions and the dotted arrows to the backward reactions. The upward facing arrows indicate where photon absorption (> 4 eV, UV-C) is required for the reaction to proceed. Image credit: Boulanger et al. (2013).

The full reaction requires the photo-excitation of cis-DAMN (1), trans-DAMN (4), and AIAC (5), with photons of over 4.0 eV (UV-C). Koch and Rodehorst (1974) have found that irradiation with UV-C light induces photoisomerization of cis-DAMN, leading to a photostationary state with a large predominance of trans-DAMN (4) isomer over cis-DAMN (1). This cis-trans isomerization under the UV-C light involves three internal coordinates; rotation around the C=C bond, the pyramidalization of one of the carbon atoms which stabilizes charge transfer in the photon-induced ^1^ππ* transition (Levine and Martinez, 2007) and the elongation of the C-N bond (see figure 6). There are also two main types of cis–trans photoisomerization mechanisms through either singlet or triplet states (Waldeck, 1991), each being competitive in the of cis-DAMN (1) to trans-DAMN (4) isomerization (Szabla et al., 2014). The measure of the θ angle (figure 6) amounts to 3.8 and 181.98 ° for cis-DAMN and trans-DAMN, respectively (Szabla et al., 2014). At the MP2 level of optimization, Cis-DAMN is the thermodynamically more stable isomer (ΔG_cis–trans_=0.61 kcal Mol_-1_) owing to less strained steric interaction in the ground state (Szabla et al., 2014).

**Fig. 6:**
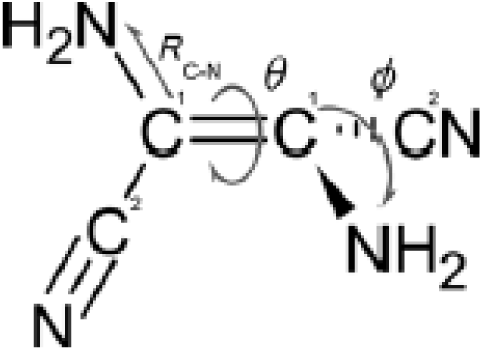
There are in general three internal coordinates involved in the photoisomerization of cis- to trans-DAMN. The first is the rotation angle θ about the C=C double bond, and the second is the pyramidalization coordinate ϕ of the C1’ carbon atom defined as the angle between the C-N bond and the plane defined by the C1=C1’-C2’ plane, and the third is the elongation of the C-N bond. Image credit: Szabla et al. (2014).

The wavelength of the ^1^ππ* transition was predicted by Szabla et al. (2014) to be 264 and 300 nm for cis-DAMN and trans-DAMN, respectively, with a corresponding red shift when the calculation was carried out in a polar solvent. This non-equilibrium isomerization is a clear example of microscopic dissipative structuring involving a continuous photon flux and the internal electronic and isomeric degrees of freedom of the molecule.

An important result found by Boulanger et al. (2013) is that although energies of greater than 4.0 eV are added to the system through photoexcitation by UV-C photons, most of this free energy, rather than being consumed in the photochemical reactions, is instead simply dissipated to the ground state of the relevant structure through internal conversion occurring in less than 0.2 ps. In fact, Koch and Rodehorst (1974) have shown that cis-DAMN (Fig. 4 (1)) is excited about 300 times (on average) before a single cyclization event (Fig. 4 (3)) takes place. This photochemical formation of the purines from HCN is thus a very dissipative process and the experimental data presented by Koch and Rodehorst, along with the molecular dynamic simulations of Boulanger et al. and Szabla et al., strongly suggest that individual steps in the overall observed process of purine formation from HCN under a UV-C flux are examples of microscopic dissipative structuring in which molecular structures (pigments) arise “spontaneously” to dissipate the same external generalized chemical potential that produced them.

Assuming that the nucleobases were formed during the Archean through microscopic dissipative structuring under UV-C light, as suggested for the purines in this section, it would have been crucial to the dissipative program to keep the nucleobases at the surface of the ocean where they would have been exposed to maximum UV-C light. Besides a class of fundamental amino acids which are hydrophobic, sugars also demonstrate significant hydrophobicity. Of all the common sugars, ribose is the most hydrophobic (Janado and Yano, 1985), so an association of a nucleobase with ribose, forming a nucleoside, would have been beneficial in preventing sedimentation and thereby optimizing photon dissipation. Ribose is also very stable under UV-C light, and to even shorter wavelength radiation, due to its own inherent conical intersection (Tuna et al., 2016).

## 5. Microscopic Dissipative Structuring of Single Strand RNA or DNA

The following step in the evolution of life was most probably the polymerization of the nucleosides into single strand RNA or DNA. There are both thermodynamic and non-equilibrium thermodynamic incentives for the polymerization which are, respectively; greater stability against hydrolysis, and the provision of a scaffolding to allow useful molecules to attach themselves to RNA or DNA. For example, antenna type molecules which, through Förster-type energy transfer, could have used the conical intersections of the acceptor nucleic acid molecules to rapidly dissipate their UV-C induced electronic excitation energy into heat (Michaelian, 2016), or, hydrophobic molecules which could have trapped the polymers at the surface for greater dissipation. This section shows how these thermodynamic principles of microscopic dissipative structuring could have led to the autocatalytic formation of single strand RNA or DNA given the existence of nucleobases and a UV-C light flux in the environment.

Consider the following example of a simplified reaction scheme for the template directed autocatalytic photochemical production of single strand DNA or RNA polymers of length 10 nucleotides. Experimental evidence for the viability of this set of reactions in the context of the prevailing Archean environmental conditions, which we have called Ultraviolet and Temperature Assisted Replication (UVTAR), can be found in Michaelian and Santillán (2014) and Michaelian (2016). The reactions to be considered are;

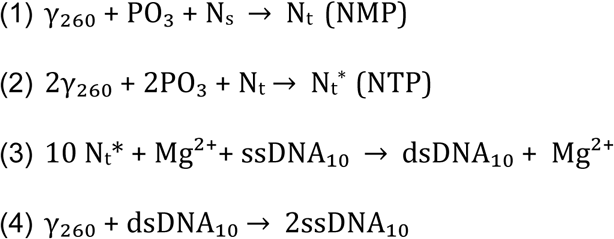

In reaction (1), a 260 nm UV-C photon interacts with a phosphate group PO3 close to a nucleoside N_s_ to produce a nucleotide N_t._ (nucleotide monophosphate). In reaction (2), two similar photons interact consecutively with two phosphate groups to produce an activated (phos-phorylated) nucleotide N_t_* (nucleotide triphosphate) (McReynolds et al.,1971). In the third reaction (3), 10 activated nucleotides interact, over night, at somewhat colder temperatures and with the aid of Mg^2+^ ions acting as catalysts for their polymerization (Szostak, 2012), with a single strand DNA segment of 10 nucleotides, ssDNA_10_, acting as a template to produce a double strand DNA, dsDNA_10_. In the final photochemical reaction (4), occurring during the day, a 260 nm UV-C photon interacts with a double strand DNA and, through the dissipation of the excitation energy into local heat, and the puckering of the C_2_ atom (Adenine) or the C_6_ atom (Guannine), and the out-of-plane distortion of the NH_2_ groups of the excited base (adenine, guanine, and cytosine) needed to reach the conical intersection (Natchtigallova et al., 2010, Plasser et al., 2014), hydrogen bonds are broken and the double helix denatures it into two single strands, ssDNA_10_ (Michaelian 2011, 2016). The breaking of hydrogen bonds could also be initiated by the formation of photon-induced OH_-_ radicals causing deprotonation of DNA.

Note that the viability of each of the above four reactions has been demonstrated independently experimentally and there does not appear to be any in principle reason why the set of reactions could not occur in the given sequence under Archean ambient conditions (Michaelian, 2016). Note also that because of the nature of these reactions involving high energy photons, the backward rate constants would be essentially zero, implying the reaction set is far from equilibrium.

The overall photochemical reaction can thus be written as;

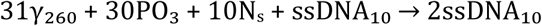

which is just an autocatalytic template directed photochemical reaction of the form A + B →2B for the production of single strand DNA, with A = 31_γ260_ + 30PO_3_ + 10N_s_ and B = ssDNA_10_.

## 6. Dissipative Proliferation of the Fundamental Molecules

Following Prigogine's non-equilibrium thermodynamic analysis of autocatalytic chemical reactions (Prigogine, 1967), I have shown (Michaelian, 2013) that, if under the imposition of the solar photon potential, and given a constant supply of reactants and a constant sink (dispersal) of the products, a photochemical route can be found to the production of a pigment molecule (such as the two examples given above), and if that pigment molecule is efficient at dissipating the same photon potential that was required to produce it, then, a process similar to that of the autocatalytic chemical reaction would occur, only that a photochemical potential, rather than a chemical potential, would be dissipated. In a manner analogous to what happens with the product catalyst in a chemical autocatalytic reaction, the concentration of the pigment in the photochemical autocatalytic reaction at the thermodynamic stationary state (when the rates of all dissipative processes are no longer changing, given a constant imposed external photon potential and a constant source of reactants and sink of products) would become many orders of magnitude larger than what would be expected under near equilibrium conditions. The derivation shows that the greater the efficacy of the particular pigment in dissipating the solar photon potential, the greater the amount of that pigment in the stationary state that would be expected and the greater the global entropy production of the process in the stationary state (Michaelian, 2013). This is just the replication-dissipation relation alluded to above which is responsible for life’s perceived “vitality”.

The above mechanisms of UV-C induced isomerization, activation of nucleotides, and denaturing in autocatalytic photochemical production of single strand DNA, are not the only mechanisms available for dissipative structuring. As another example, consider the situation in which a UV-C photon is capable of converting a molecule A plus a molecule B into a pigment molecule C. Photochemical reactions are like thermal reactions in the sense that higher temperatures imply more rapid and extensive movement in configuration space (the coordinate space of the atoms of the molecules) which is helpful in finding the optimal collision orientation between molecules A and B for the reaction to take place. Rate constants for both chemical and photochemical reactions are thus normally increasing functions of the local solvent temperature. If the pigments C produced in a photochemical reaction are effective converting the photon energy rapidly into local heat, their presence could increase the rate of the photochemical reaction producing the pigment itself, thereby acting as a catalyst for its own production.

Not only could the pigment provide local heat, but it may also be able to transfer energy in other ways, such as through Förster resonant energy transfer, through charge transfer, or fluorescence, etc. In such autocatalytic reactions, the concentration of pigments C will increase to a value at the stationary state many orders of magnitude larger than what it would have been if the pigment did not have its catalytic properties, or if the reaction were close to equilibrium. An inexhaustible source of UV-C photons and primordial molecules (HCN) and dispersal (sink) of the newly formed pigment catalyst, by whatever mechanism, will thus imply a continual local production of the pigment to maintain the stationary state, a process which could be coined as “dissipative proliferation”.

## 7. Evolution

Whatever molecular association or novelty that increased the entropy production of the fundamental molecules when exposed to a UV-C flux would therefore be proliferated (differentially selected) through this dissipation-replication relation. As an example, the 3-base pair stereochemical coding in RNA or DNA for tryptophan (Yarus et al., 2009), another UV-C pigment (see figure 3), may have happened because tryptophan could have “utilized” (in a non-equilibrium thermodynamic sense) the conical intersection of RNA or DNA to rapidly dissipate its UV-C photon induced electronic excitation energy, the complex dissipating more efficiently than the component molecules acting independently. The RNA or DNA codon specifying for an antenna molecule such as tryptophan could thus be considered as part of a compound microscopic dissipative structure created during the Archean that has persisted, and is even utilized, to this day, even though the external generalized chemical potential that created and proliferated it (the UV-C photon potential) no longer exists at Earth’s surface. RNA and DNA thus eventually took on the new role of information storage, with the stored information invariably associated with the dissipation of an externally imposed generalized chemical potential at one time existent in the organisms environment (Mejía and Michaelian, 2017).

It is thus possible to envision increases in complexity concomitant to increases in dissipation and nested sets of autocatalytic photochemical and chemical reactions occurring, each feeding off a more fundamental photochemical reaction process by using its products and the generalized chemical potential generated (e.g. heat gradients or chemical potential). An instructive example of precisely this was given above in the photochemical production, starting from HCN, of single strand RNA or DNA. This is similar to Kaufmann’s (1986) suggestion for the origin of life resulting from the emergence of a set of autocatalytic chemical reactions, but where we have identified an external UV-C photon potential as the fundamental driver of the photochemical dissipative structuring and proliferation of the products (fundamental molecules) for the dissipation of this potential.

For such sets of autocatalytic photochemical and chemical reactions, the overall coupled process generates more entropy than could the individual processes summed (if indeed they could be artificially separated). This hierarchy of autocatalytic dissipative cycles is the scheme employed ubiquitously by life today. Examples are the metabolic Krebs cycle and the carbon fixation in the photosynthetic cycle.

It is the dissipation of the solar photon potential by these fundamental molecules of life (pigments in the UV-C), through the type of autocatalytic photochemical reactions detailed above, that was, and still is, driving their proliferation over the sunlit Earth. This photon driven proliferation we identify at macroscopic scales as the “vitality” of life. These molecules, and their complexes forming greater dissipating structures (from molecular pigment complexes to the biosphere) do not have a mysterious "will to survive and proliferate" as the traditional evolutionists are obliged to contend, rather, they are driven to proliferation based on how good a catalyst they are for dissipating the external photon potential. Recognizing this fact provides a non-tautological basis for understanding evolution within a thermodynamic framework based on established physical and chemical law.

Today, the Earth's biosphere dissipates into heat a large number of solar photons arriving at Earth's surface by utilizing an enormous quantity and variety of organic pigments, of which chlorophyll is just one notable case. For example, the carotenoids are known acceptor quencher molecules for the rapid dissipation (within 200 ps) of excited chlorophyll donor molecules, the two pigments being held close enough together for energy transfer in high-light inducible proteins (Hilps) which are the cyanobacterial predecessors of the light harvesting complexes (LHC) found in plants (Staleva et al., 2015). The carotenoid-chlorophyll dissipative system may thus be considered a modern analog of the RNA/DNA-tryptophan dissipative system of the Archean discussed above.

Even with such a formidable array of evolved pigment complexes, the wavelength integrated flux of photons from the Sun at Earth's surface is so copious (∼10^22^ m^−2^s^−1^) that only a portion of the solar photons arriving at Earth's surface are absorbed and dissipated by today's evolved pigments in plants, diatoms and cyanobacteria (Michaelian and Simeonov, 2015). The rest of the solar photons, principally those with wavelength longer than that corresponding to the red-edge (∼700 nm), are simply reflected with practically no energy dissipation. However, even this diffuse reflection produces entropy by distributing photon energy over a much larger solid angle of space than that corresponding to the incident beam (Michaelian, 2012).

The thermodynamically most important co-evolutionary (biotic-abiotic) process occurring with the biosphere has been that of increasing the photon dissipation efficiency of Earth in its solar environment and this evolution has continued ever since the formation of Earth in the Hadean until the present era. There seems to be no better or succinct description of biotic-abiotic co-evolution than this observation. The evolution of ever greater photon dissipation efficacy of the biosphere has incurred all of the following: Evolving,

1. a greater photon absorption cross section per pigment size, for example by increasing molecular electric dipole moments,
2. faster non-radiative dissipation of the photon-induced electronic excitation energy of the pigment through the evolution of molecular conical intersections so that the complex promptly returns to the ground state, ready to process another photon,
3. quenching of the fluorescent and phosphorescent radiative decay channels, through, for example, developing structures which promote resonant energy transfer,
4. quenching of electron transfer reactions, except where required for specific dissipative structuring processes, e.g. nucleobase structuring and photosynthesis,
5. stability against photochemical reactions and electron transfer reactions which could destroy the molecule,
6. new pigments absorbing at different wavelengths by extending conjugation in a molecule, thereby covering ever more completely the solar spectrum at Earth’s surface,
7. processes that exude pigments into the environment to promote dissipation over a greater part of the solar spectrum, for example, oxygenic photosynthesis leading to oxygen and ozone in the environment,
8. mechanisms for promoting the dispersal of pigments over ever more of Earth's sunlit surface and into new inhospitable environments, through, for example, the evolution of the complex cell, insects and other animals required for dispersing nutrients and seeds; a process fomented by the exudation into the environment of the pigment oxygen (point 7),
9. greater coupling of photon dissipation in organic pigments to other biotic and even abiotic irreversible processes, e.g. the water cycle, winds and ocean currents,
10. transparency of Earth's atmosphere to the most intense high energy part of the Sun's spectrum so that photons can arrive at Earth’s surface where they can be best intercepted by complex organic pigments in liquid water and thereby dissipated most efficiently.

The complexification of material into mobile abiotic or biotic mechanisms (organisms) for transporting the pigments (and the nutrients required for their proliferation) into regions where sunlight exists but where nutrient content is low (point 8) provides a thermodynamic explanation for the proliferation of the water cycle and animals. Humans appear to have had an important role in fomenting the amount of carbon in the carbon-cycle and water in the water-cycle, and the future may involve an extension of this thermodynamic function to other planets, and even an alternative role; that of diverting sunlight into regions of high organic nutrient content, and this re-direction of generalized thermodynamic forces may be a useful thermodynamic definition of an "intelligent" animal.

## 8. Summary

The physical, chemical, optical, and electronic properties of RNA and DNA and the other fundamental molecules of life suggest that these molecules were once microscopic dissipative structures that originated and proliferated over the ocean surface as an irreversible thermodynamic imperative to absorb and dissipate the prevailing Archean solar UV-C photon flux. Their proliferation to far beyond concentrations expected under near equilibrium conditions can be understood in terms of non-linear, non-equilibrium thermodynamic principles directing autocatalytic photochemical reactions in which these pigments catalyze the dissipation of the same thermodynamic potential (the solar photon flux) that produces them. The non-equilibrium thermodynamic framework presented here is in line with Boltzmann's (1886) extraordinary insight into the nature of the Darwinian struggle for existence in terms of entropy production and with Prigogine's formal analysis of spontaneous non-equilibrium dissipative structuring of material under a generalized chemical potential such that the potential is dissipated most efficiently.

It is likely that the nucleobases were originally derived from HCN through microscopic dissipative structuring under a constant UV-C photon flux through the autocatalytic photochemical reactions first studied by Ferris and Orgel (1966). I have further suggested an autocatalytic photochemical reaction set leading to an exponential increase over time in particular single strand RNA or DNA oligos that dissipated strongly the Archean UV-C flux, which we have named UltraViolet and Temperature Assisted Replication (UVTAR).

Complexification of life would have been driven by differential proliferation of complexes which increase photon dissipation efficacy most through this replication-dissipation relation. For example, by utilizing antenna UV-C pigment molecules, Förster energy transfer, and the extremely rapid dissipative characteristics of RNA and DNA afforded by inherent conical intersections. In fact, any innovation that improved photon dissipation, such as stereochemical coding for either antenna molecules or hydrophobic molecules (allowing the complex to adhere to the ocean surface where the UV-C flux was greatest), would have been thermodynamically selected since replication (proliferation) would have been tied to dissipation through the UVTAR mechanism.

Photon dissipation continues to this day as the most important thermodynamic work performed by life, but now with pigments operating in the near UV and visible regions of the solar spectrum where the solar light is most intense, while dissipation of the UV-B and UV-C regions is mostly performed by ozone derived from oxygen (which should also be considered as a pigment) exuded by photosynthetic organisms into the environment. The production of pigments today to far beyond their expected equilibrium concentrations is similarly driven, but now indirectly, through complex biosynthetic pathways employing autocatalytic photochemical reactions operating in the near UV and visible.

Evolution has been, and still is, primarily concerned with increasing the rate of photon dissipation in the biosphere and this has been achieved by evolving pigments with extraordinarily rapid times for conversion of electronic excitation energy into vibrational energy, evolving pigments of ever larger photon cross section and of increased spectral range, more rapid non-radiative decay rates while at the same time reducing the probability of molecular destruction or radiative decay (e.g. fluorescence and phosphorescence), and proliferating these pigments with the aid of the water cycle and mobile animals which helped them to spread over ever more of Earth's surface, thereby increasing global photon dissipation.

The implicit, but unjustified, assumption of an inherent organismal "will to survive" in the Darwinian theory can now be replaced with an explicit and physically founded "will to produce entropy" (colloquially speaking) in the thermodynamic dissipation theory of the origin and evolution of life. However, we should no longer speak about individual selection (although this still retains a residual meaning if put in terms of an individual’s contribution to global entropy production instead of fitness) but instead about hierarchal thermodynamic selection starting with the fundamental molecules (nucleobases, RNA/DNA, aromatic amino acids, and other fundamental molecules which are UV-C pigments) up to the biosphere including the UV and visible pigments and involving both its biotic and abiotic components. Random associations and perturbations occur simultaneously at all levels, including at the individual organismal level, biosphere level, and even at the most microscopic pigment molecular levels, and these complexes are "selected" (a better word would be "proliferated") based on how good a catalytic agent they are in fomenting the global entropy production given both local and global constraints (photon potential, nutrients, water, etc.). What is optimized in nature is not organismal "fitness", which, in fact, cannot even vaguely be defined without incurring in circular arguments, but rather "global entropy production" which can, and, in fact, has been, accurately measured (Michaelian, 2012). A remaining challenge is to reframe all of the accepted postulates of Darwinian Theory from within this non-equilibrium thermodynamic framework.

Microscopic self-organization through dissipation and the persistence of nanoscale structuring at the origin of life would not be unique to Earth but should have occurred wherever there once existed UV-C or UV-B light and organic elements; on protoplanetary systems, on planets, on asteroids, on comets, and even in the galactic gas and dust clouds (Michaelian and Simeonov, 2017). Organic pigments on Titan catalyzing the methane rain cycle, or the sulfur containing pigments floating in the clouds of Venus catalyzing the great southern and northern vortices, are examples of microscopic dissipative structuring giving rise to “living” biospheres on other planets. The cosmos should be teaming with such biotic-abiotic ecosystems based on pigments “self-organized” through microscopic dissipative structuring under the photon potential of the local star.

## Acknowledgements

The author is grateful for useful suggestions on the manuscript by Carlos Bunge, and for financial support granted by the Dirección General de Asuntos del Personel Academico, DGAPA-UNAM, project number IN102316.

